# Deficiency of IL-36 receptor antagonist (DITRA) is associated with decreased homoeostatic CCL27 expression leading to heightened dermal inflammation

**DOI:** 10.64898/2026.06.11.731561

**Authors:** Shrikanth Chomanahalli Basavarajappa, Paloma Narros-Fernández, Heather Loughnane, Larissa Bless, Hernandez-Santana Yasmina E, Eirini Giannoudaki, Anne C. Moore, Margaret B. Lucitt, Darren Ruane, Patrick T. Walsh

## Abstract

Deficiency of the Interleukin-36 Receptor antagonist (DITRA) is a rare autoinflammatory condition which commonly manifests as severe, recurrent episodes of Generalized Pustular Psoriasis (GPP). Loss-of-function mutations in the *IL36RN* gene result in unopposed IL-36 cytokine signalling leading to severe psoriatic inflammation, which can be successfully treated with Anti-IL-36 receptor (IL-36R) monoclonal antibodies. Despite such advances, there remain some key questions concerning how loss of a functional IL-36R antagonist predisposes to GPP, including identifying the potential impacts of *IL36RN* mutations on skin homeostasis. To address this question, we investigated the consequences of IL-36Ra deficiency using *Il36rn*^-/-^ mice, which recapitulate the severe psoriatic inflammation observed in DITRA patients. Here, we demonstrate, that in overtly healthy *Il36rn*^-/-^ mice, prior to disease onset, there is disrupted dermal immune homeostasis, characterised by decreased expression of the chemokine CCL27. Altered skin homeostasis occurred in association with dysbiosis of the skin microbiome, characterised by a significant outgrowth of the commensal bacteria, *Cutibacterium acnes*. Importantly, intradermal administration of recombinant CCL27, prior to disease induction, significantly reduced the enhanced severity of psoriasiform inflammation, demonstrating a central role for this chemokine in regulating predisposition to increased severity. Transcriptomic analysis of GPP patient’s skin also revealed decreased *CCL27* expression in non-lesional, as well as lesional, compared to healthy skin, indicating that this chemokine may also play a key instructive role among DITRA patients. Together, these data identify a novel mechanism through which IL-36Ra deficiency alters dermal homeostasis and predisposes to increased severity of psoriatic disease observed in DITRA patients.

## Main Text

Interleukin-36 receptor antagonist (IL36Ra) deficiency, caused by mutations in *IL36RN* gene, leads to an autoinflammatory syndrome in humans known as Deficiency of Interleukin-36 receptor antagonist (DITRA). DITRA is rare monogenic disease characterized by episodes of severe eruptive skin inflammation, referred to as Generalized Postural Psoriasis (GPP), orchestrated by dysregulated activity of IL-36 family cytokines. The IL-36 family of cytokines consists of three distinct ligands, IL-36α, IL-36β and IL-36γ, which mediate inflammation in the skin through engagement with a distinct Interleukin-36 receptor (IL-36R). Under homeostatic conditions, this axis is tightly regulated through the expression and activity of IL-36Ra, which can bind to IL-36R with higher affinity and restrict cytokine signalling ^1^. Accordingly, loss-of-function mutations in *IL36RN* disrupt this regulation, predisposing to the development of GPP ^2^.

While a deeper understanding of the mechanisms which drive the pathogenesis of DITRA led to the recent approval of spesolimab (Anti-IL-36R) as a first-in-class therapeutic for the treatment of GPP ^3^, several questions remain with respect to how IL-36Ra loss of function predisposes to disease onset. The age of onset among DITRA patients is known to vary widely, ranging from early infancy to middle age ^4^, highlighting the role of poorly defined environmental triggers to initiate disease, while also indicating a potential dysregulation of healthy dermal homeostasis among DITRA patients. To address these questions, we sought to investigate whether deficiency of the IL-36Ra resulted in alterations in the dermal immune barrier interface under homeostatic conditions. While *Il36rn*^*-/-*^ mice exhibit increased severity of Aldara-induced psoriasiform inflammation (sFig.1A-C), they are otherwise overtly normal and do not display any evidence of dysregulated skin barrier function or inflammation when raised under specific pathogen-free conditions. Therefore, we chose these mice as a relevant model to determine whether loss of IL-36Ra function might alter dermal homeostasis and predispose to increased severity of psoriatic inflammation.

An analysis of altered gene expression profiles, through RNA sequencing, in *Il36rn*^*-/-*^ skin revealed limited significant alterations but a notable decrease in levels of the chemokine *Ccl27*, an established mediator of skin immune homeostasis (Fig. 1A). CCL27, which is predominantly expressed by keratinocytes, has been described to promote dermal homeostasis through the recruitment of CCR10^+^ immune cells such as T cell subsets, including CD4^+^FoxP3^+^ regulatory T cells (Treg), as well as innate lymphoid cell (ILC) subsets to the skin ^5,6^. While, CCL27 expression has previously been shown to be decreased in lesional skin from psoriasis patients, its role in GPP pathogenesis is yet to be described. Decreased CCL27 protein expression was evident both in the skin and serum, as well as in primary keratinocytes derived from healthy *Il36rn*^*-/-*^ mice (Fig. 1B-D). This decrease in expression also occurred in association with reduced numbers of skin resident ILC (Fig.1E-G), while CD4^+^FoxP3^+^ Tregs were unchanged (sFig.2C-D), demonstrating a dysregulated dermal immune interface in absence of IL-36Ra activity. Interestingly, these alterations were also found to occur in association with dysbiosis of the skin microbiome characterised by a specific outgrowth of the skin commensal bacteria, *Cutibacterium acnes (C.acnes)*, further demonstrating a homeostatic role for constitutive IL-36Ra expression in the skin (Fig. 1H).

**Figure 1:**
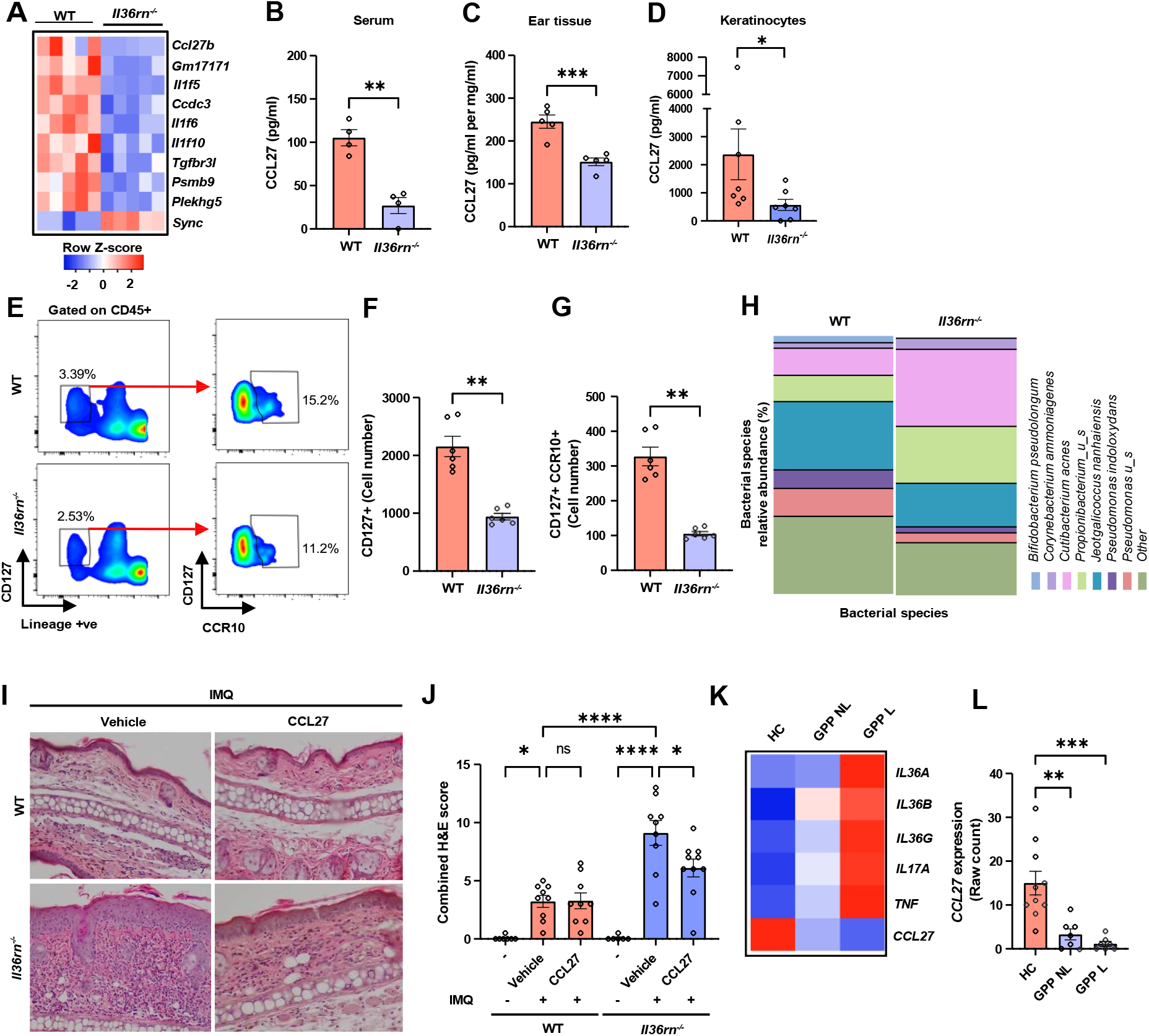
Deficiency of IL-36R antagonist results in reduced CCL27 expression and altered skin homeostasis. (A) Heatmap illustrating significant differentially expressed genes in *Il36rn*^*-/-*^ skin compared to wild type littermate controls (WT)(P.adj<0.05; n=5 mice per group). (B-D) Analysis of CCL27 protein levels in serum (n=4), ear skin homogenates (n=5) and primary keratinocytes (n=6) of WT and *Il36rn*^*-/-*^ groups as determined by ELISA. (E-G) Representative FACS plots (left) and corresponding bar graphs (right) showing the percentage and relative cell numbers of CD45^+^Lin^-^CD127^+^ cells and CD45^+^Lin^-^CD127^+^CCR10^+^ cells in WT and *Il36rn*^*-/-*^ mice skin. n=6 mice per group. (H) Heatmap demonstrating relative abundance of identified bacterial species as determined by shotgun metagenomic sequencing in WT (n=5) and *Il36rn*^*-/-*^ (n=5) mice.(I) Administration of CCL27 reversed increased severity of psoriasiform inflammation in *Il36rn*^*-/-*^ mice as determined by by H&E staining (J) and combined histological score of ear sections from the indicated groups on day 6. (I & J) Combined data from two independent experiments. (K) Analysis of publicly available bulk RNA sequencing dataset E-MTAB-11144 showing a heatmap summarizing the average expression of key inflammatory cytokines implicated in GPP pathogenesis, along with *CCL27*. (L) *CCL27* expression in non-lesional and lesional skin of GPP patients (healthy controls (HC) (n=10) and GPP patients (n=7)). Statistical comparisons in (B-D & L) were performed using unpaired t test: ** p<0.01,*** P<0.001,in (F & G) were performed using Mann-Whitney test: ** p<0.01, and in (J) were performed using ordinary one-way ANOVA multiple-comparison test: ns p>0.05, * p<0.05, **** P<0.0001.

Importantly, the consequences of decreased CCL27 expression in *Il36rn*^*-/-*^ mice, with respect to the pathogenesis of more severe psoriasiform disease, was demonstrated through a significant reduction in the severity of Aldara-induced psoriatic inflammation, upon intradermal administration of recombinant CCL27 (Fig.1I & J). Moreover, analysis of skin biopsy gene expression from a cohort of GPP patients demonstrated that *CCL27* expression levels are significantly decreased in non-inflamed, as well as inflamed skin (Fig.1K & L). As well as mirroring our murine data, this suggests that CCL27 activity may play an important regulatory axis in GPP pathogenesis.

Mechanistically, stimulation of primary keratinocytes with recombinant IL-17A decreased CCL27 secretion (sFig.3A), whereas reduced *Ccl27 and Ccr10 gene* expression in the skin of wild type mice, induced by Aldara treatment, was found to be reversed upon IL-36R or IL-12/23 blockade *in vivo* (sFig.3B). In addition, *CCL27* expression was inversely correlated with *IL36* family and *IL17A* gene expression in GPP patient’s skin (sFig.3C-F). While these data indicate a role for IL-17A in regulating CCL27 expression and activity at the skin barrier, whether this is occurs under homeostatic conditions, or is linked to microbial dysbiosis on the skin surface, remains to be determined.

These data provide the first evidence that the skin homeostatic chemokine CCL27 plays a central role in the pathogenesis of DITRA and offers novel insights into the mechanisms underlying this rare autoinflammatory disease. Furthermore, these data suggest that CCL27-based therapeutics could help to mitigate the pathogenesis of GPP among DITRA patients.

## Funding Support

PTW acknowledges funding support from Science Foundation Ireland (21/FFP-P/10135), and the Higher Education Authority (HEA) North-South Research Programme; “All-island Vaccine Research and Training Alliance (AIVRT)” to ACM and PTW.

## Notes

**Competing Interest Statement:** The authors declare that they have no conflict of interest.

### Competing Interest Statement

The authors have declared no competing interest.

