## Supplemental Methods and Figures 1-3 for "Deficiency of IL-36 receptor antagonist (DITRA) is associated with decreased homoeostatic CCL27 expression leading to heightened dermal inflammation"

**Supplementary Material:**

**Methods:**

**Mice:**

Wild-type (WT, C57BL/6J) littermate controls (LMC) and *Il36rn*$^{-/-}$ mice (provided by M. Kopf, University of Zurich) were bred in-house under specific pathogen-free conditions at the Trinity Translational Medicine Institute, Dublin, Ireland. For all experiments, both male and female mice aged 8-12 week were used. For microbiome studies and skin RNA sequencing studies, *Il36rn*$^{-/-}$ mice and WT LMC were cohoused from birth in the same cages until analysis at 12 weeks.

**Sex as a biological variable:**

Both male and female mice were used in this study and similar results were observed for both sexes.

**Primary culture and treatments:**

Primary keratinocytes were isolated from the skin of WT and *Il36rn*$^{-/-}$ mice skin as described earlier. Cells were stimulated with IL-17a (100 ng/ml; #421-ML, R&D systems) for 24 hours. Following stimulation, supernatant and cell pellet were collected and stored at -80ºC for further use.

**DNA sequencing and analysis of skin microbiome:**

Skin microbiome samples from cohoused WT LMC and *Il36rn*$^{-/-}$ mice were collected using sterile swab pre-moistened with sterile PBS containing Tween20 (0.05%). Swabs were gently rolled over the shaved back skin for 2 min and immediately frozen for further processing. Frozen swabs were then sent to CosmosID for DNA sequencing analysis. DNA libraries were prepared using the Nextera XT DNA Library

Preparation Kit (Illumina) and IDT Unique Dual Indexes, with a total DNA input of 1ng. Genomic DNA was fragmented using a proportional amount of Illumina Nextera XT fragmentation enzyme. Unique dual indexes were added to each sample, followed by 12 cycles of PCR to construct libraries. DNA libraries were purified using AMpure magnetic Beads (Beckman Coulter) and eluted in QIAGEN EB buffer. DNA libraries were quantified using Qubit 4 fluorometer and Qubit™ dsDNA HS Assay Kit. Libraries were sequenced on an Illumina HiSeq X platform with paired-end reads of 2x150bp. Bioinformatic analysis was performed by CosmosID using reference genomes database, virulence markers, and antimicrobial resistance markers. Relative abundance estimates for the microbial reads were given by CosmosID to compare the skin microbiome profile of WT and *Il36rn*$^{-/-}$mice.

**RNA extraction**

Ear tissue was lysed in RNA lysis buffer using Teen Prep Lysing Matrix tubes (#11412420, MP biomedicals) and the FastPrep-24 5G system (MP Biomedicals). Total RNA was isolated from tissue lysate using ISOLATE II RNA Mini Kit (#BIO52072, Meridian bioscience) according to the manufacture instructions and stored in -80ºC for further use.

**RNA sequencing.**

Total RNA from ear tissue was extracted as described previously and sent to Novogene (UK) for further processing. Poly-A enrichment was performed to isolate mRNA and prepare mRNA libraries from the total RNA samples. Libraries were sequenced on a Ilumina NovaSeq PE150 platform using a paired-end 150 bp sequencing strategy (short-reads). Bioinformatic analysis, including gene expression and functional analysis, was conducted by Novogene (UK).

**ELISA**

Ear tissue and cell pellets were homogenized in RIPA buffer containing protease inhibitor cocktail (#P8340, Sigma-Aldrich). Ear tissue homogenization was performed using Teen Prep Lysing Matrix tubes (#11412420, MP biomedicals) and FastPrep-24 5G system (MP Biomedicals). The total Protein concentration in the ear homogenate was determined using the BCA protein assay kit (#BCA1, Sigma-Aldrich). Cell culture samples (supernatant and cell homogenate), *Il36rn*$^{-/-}$ serum, and ear tissue homogenates were assessed by ELISA for CCL27 using CCL27/CTAK Duoset ELISA kit (#DY725, R&D Systems) according to the manufacture instructions. Protein levels in the ear and cell homogenates were normalised to the total protein concentration.

**Flow cytometry**

Ears tissue was digested with preheated HBSS (37ºC) (#14170-088; ThermoFisher scientific) supplemented with collagenase type IV (1mg/mL; #C4-BIOC, Sigma-Aldrich) and DNase I (100ug/mL; #10104159001, Sigma-Aldrich) for 90 min at 37ºC using a shaker incubator. The digested ear tissue was homogenized using syringe and passed though 100 µm and 40 µm cell strainer. Samples were then centrifuged at 1700 RPM for 7 min at 4ºC, and the resultant pellet was resuspended in FACS buffer. After cell counting, cells from 5-6 mice were pooled in each group and used for cell staining. 4 x $10^6$ cells were blocked with anti-mouse CD16/CD32 (#14-0161-85, ThermoFisher scientific) for 10 min at 4ºC, followed by surface staining using appropriate antibodies for 45 min at 4ºC. For intracellular staining, cells were first fixed and permeabilized using FOXP3 Fix/Perm Buffer set (#421403, BioLegend) and then stained overnight with appropriate antibodies at 4ºC. Stained cells were analysed by BD LSRFortessa flow cytometer (BD Biosciences). Splenocytes were used for fluorophore compensation, and data analysis was performed using FlowJo software. Gating strategies and FMO controls are shown in sFig.2A & B

**Induction of Aldara (IMQ) induced psoriatic inflammation**

Male and female twelve-week-old mice were randomly assigned to either the vehicle or CCL27-treated group. Treatments and experimental duration are outlined in Fig. 5A. To induce psoriatic inflammation, a topical application of Aldara cream (5% IMQ; MEDA Pharmaceuticals) or control cream (Vaseline) was carried out on different ears of the same mice in both WT or *Il36rn*$^{-/-}$ (26). Vehicle (0.1% PBS-BSA) or 2 µg of Recombinant mouse CCL27/CTACK protein (#725-CK, R&D Systems) was delivered via intradermal injection (20 µl) to the IMQ treated ears on day 0, 3, and 5. Ear thickness was measured daily using digital calipers (Hitec). The dose of recombinant mouse CCL27/CTACK protein used was previously described (1). Mice were euthanized at indicated timepoints, and ears were collected and fixed in 10% neutral buffered formalin (Sigma-Aldrich) for subsequent analysis.

**Histology and staining**

Ear tissues fixed in 10% neutral buffered formalin were embedded in paraffin blocks. 5-µm thick sections were prepared and stained with haematoxylin and eosin (H&E) for histological assessment. Captured images of the sections were blindly scored for acanthosis, desquamation, parakeratosis and infiltration of immune cells on a scale of 0-4 (0, no differences over control; 1, mild; 2, moderate; 3, marked; 4, severe). The Individual scores from these parameters were combined to calculate the total histological severity score .

**Analysis of public RNA sequencing data sets from DITRA mice and GPP patients**

GSE161406 (2) and E-MTAB-11144 (3) datasets were analysed for the expression of genes of interest. GSE161406 dataset contains gene expression data from RNA sequencing of humanized DITRA-like mice skin treated with PBS (n=3), IMQ (n=6), IMQ + anti-IL36-R (n=7), and IMQ + anti-IL23 (n=7). DITRA-like mice are humanized for the ectodomain of the hIL-36R and hIL-36α, hIL-36β, and hIL-36γ ligands, but not the IL-36Ra and are reported to exhibit enhanced IL-36R signalling equivalent to that observed in patients with DITRA. E-MTAB-11144 dataset includes gene expression data from the bulk RNA sequencing of skin and blood samples from individuals with General Pustular Psoriasis (GPP), Palmoplantar Pustulosis, and healthy controls (HC). Samples to be included from E-MTAB-11144 for analysis were lesional and non-lesional skin biopsies of individuals with GPP, prior to spesolimab treatment (n=7), and healthy controls (n=10). Biopsies included in analysis were from equivalent body sites (limbs). The Fragments Per Kilobase per Million mapped fragments (FPKM) in both databases were used to graph the expression patterns of genes of interest, and for correlation analysis, using GraphPad Prism 10. Differentially expressed genes (DEGs) were identified in both datasets using the Bioconductor package DESeq2 (Version 1.46.0). Genes with a $|log^2FC| > 0.5$ and FDR-adjusted p-value $< 0.05$ were considered differentially expressed. Heatmaps were generated using heatmapper.ca (29)

**Statistical analysis**

Statistical analysis were performed using GraphPad Prism 10 (GraphPad) and appropriate statistical test such as unpaired Student *t* test or Mann-Whitney or one-way ANOVA or Two-way ANOVA were used as indicated in the figure legends. For correlation analysis, Spearman correlation was used. Statistical significance was defined as $p<0.05$ (*),$p<0.01$(**),$p<0.001$(***) and $p<0.0001$(****).

**Study Approval**

All mice experiments were approved by the Trinity College Dublin Animal Research Ethics Committee and conducted under licence from the Irish Health Products Regulatory Authority (project authorization nos:AE19136/P125 and P200).

**Data Availability**

All raw data included in this manuscript is available in the Supporting Data Values file associated with this manuscript. RNA sequencing data derived from wild type and *il36rn*$^{-/-}$ mouse skin will be deposited on Gene Expression Omnibus.

**sFigure 1.**

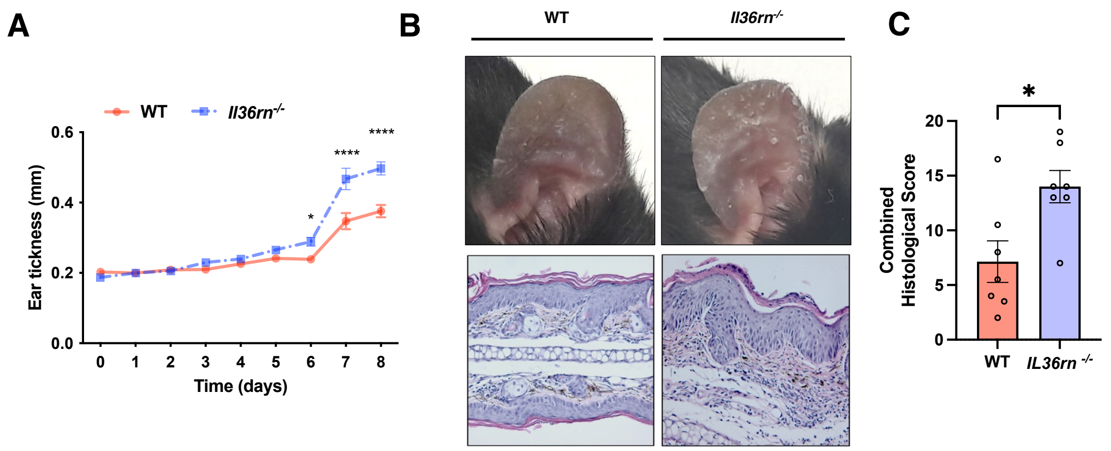


**sFigure 1: Aldara promotes psoriasiform inflammation in *Il36rn$^{-/-}$* mice.** WT and *Il36rn$^{-/-}$* mice were treated with Aldara daily and monitored for the development of psoriasiform inflammation until day 8. (A) Ear thickness from day 0 to day 8. (B) Representative clinical features of the ear skin of the mice on day 8 (top), representative H&E Staining of ear section on day 8 (bottom). (C) combined histological score on day 8. Representative data from two independent experiments (n=7 mice per group). Statistical comparison in (A) were performed using two way ANOVA Bonferroni's multiple comparisons test. Statistical comparison in (C) were performed using Mann-Whitney test: * $p<0.05$.

**sFigure 2.**

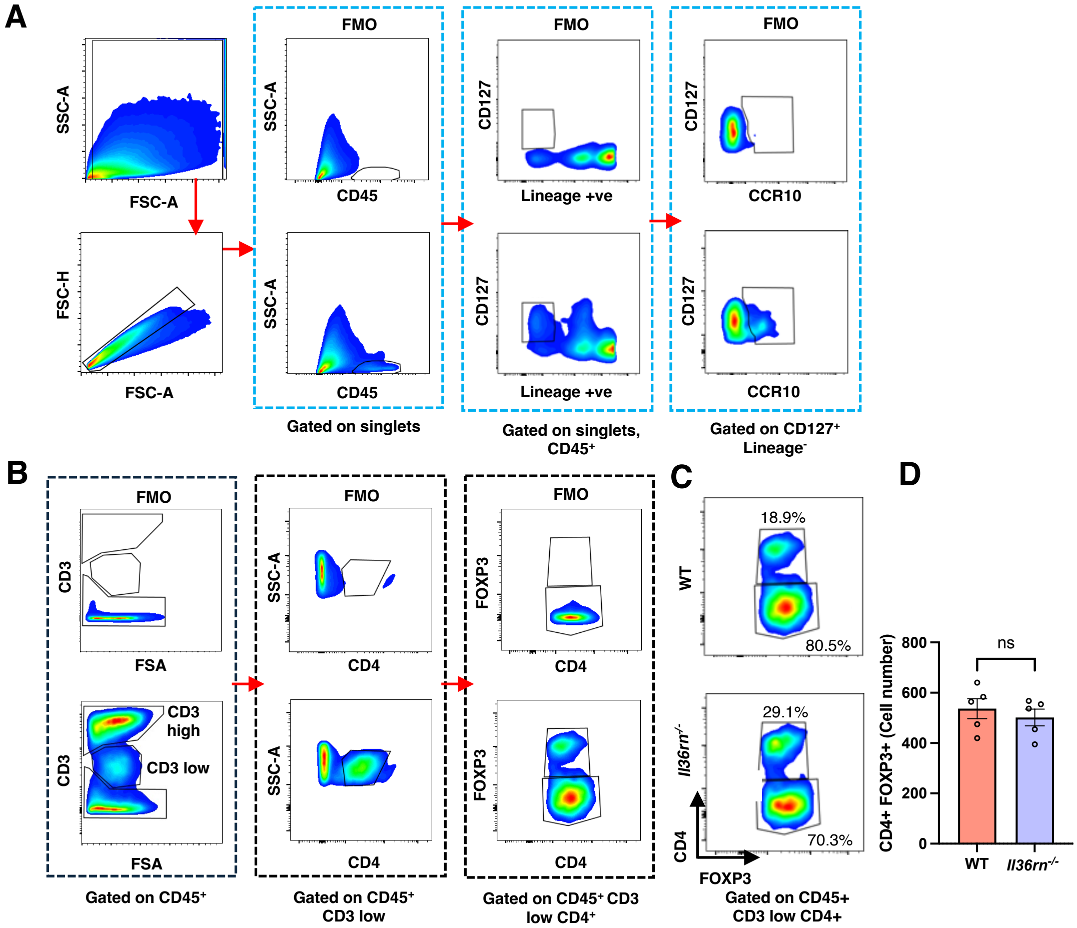


**sFigure 2: Flow cytometry analysis of ear cells from WT and *Il36rn$^{-/-}$* mice.** Flow cytometry gating strategy and fluorescence minus one (FMO) controls for the analysis of (A) CD45$^{+}$ cells, CD45$^{+}$ lineage$^{-}$ CD127$^{+}$ cells, CD45$^{+}$ lineage$^{-}$ CD127$^{+}$ CCR10$^{+}$cells (B) A similar gating strategy and FMO controls were used for the analysis of CD45$^{+}$ CD3$^{low}$ CD4$^{+}$ FOXP3$^{+}$ expressing immune cell subsets. (C & D) Representative FACS plots (left) and baer graphs (right) showing the percentage and relative cell numbers of CD4$^{+}$FoxP3$^{+}$ cells in WT and *Il36rn$^{-/-}$* skin (n=5 mice/group). Statistical comparison in D were performed using Mann-Whitney test. ns p>0.05.

**sFigure 3.**

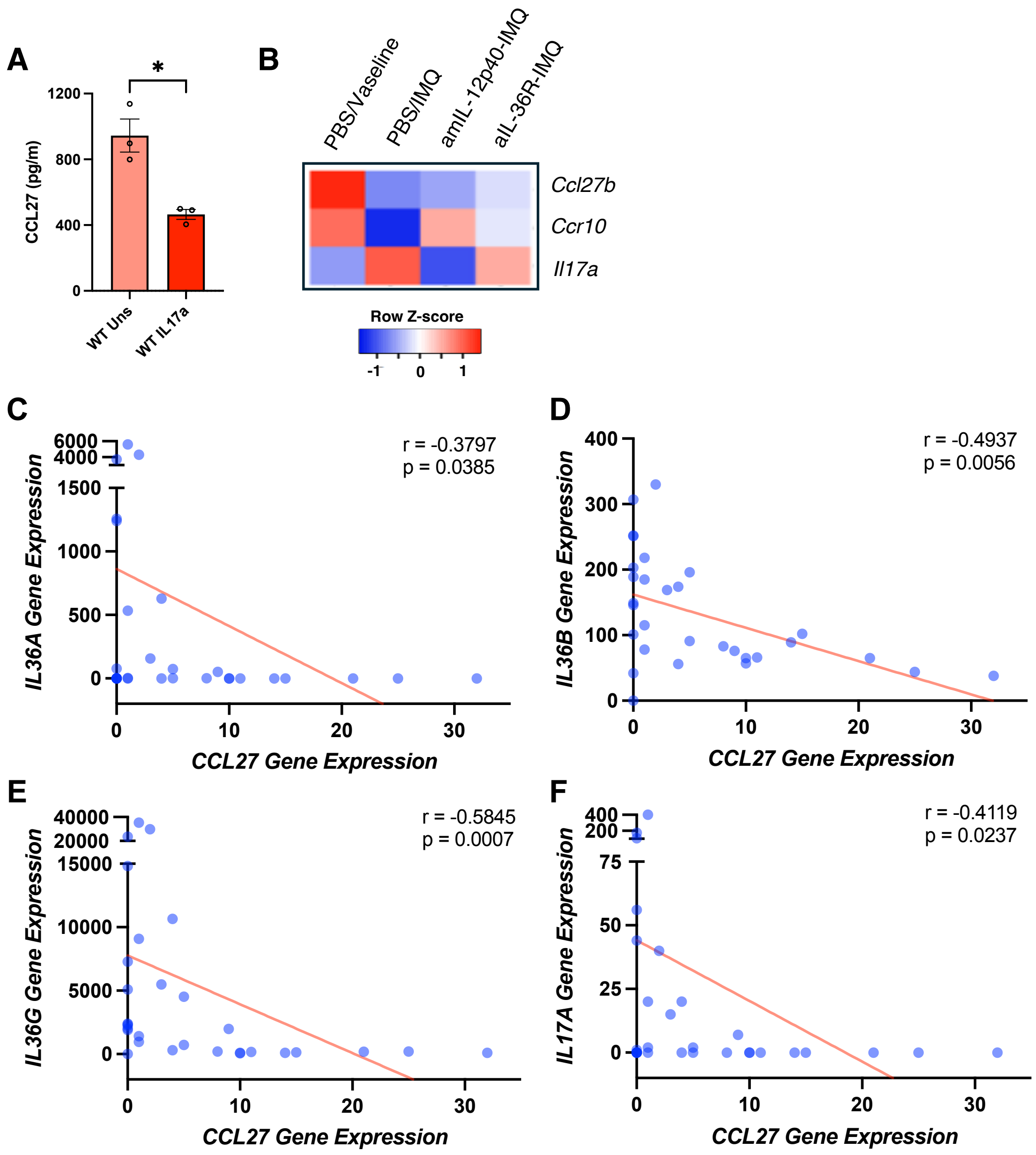


**sFigure 3: Inflammatory cytokines regulate *Ccl27* expression in keratinocytes, DITRA-like mice and correlates with *CCL27* expression in GPP pateints.** (A) Expression of CCL27 protein levels by primary keratinocytes from WT mice after treatment with IL-17a (100 ng/ml) for 24 hrs (Combined data from 3 mice). Statistical comparison performed using Mann-Whitney test: * $p<0.05$. (B) Heatmap illustrating analysed data from publicly available skin bulk RNAseq dataset GSE161406 (Gene Expression Omnibus) detailing relative expression levels of *Ccl27b*, *Ccr10*, *Il17a* across indicated treatment groups in DITRA-like mice. Relative expression derived from mean FPKM values shown as z scores. (C-F) Correlation analysis of *IL36A*, *IL36B*, *IL36G* and *IL17A* expression vs. *CCL27* expression in GPP patients (n=7) and healthy controls (HC) (n=10), derived from publicly available bulk RNA sequencing dataset E-MTAB-11144 (Biostudies). Correlation between genes of interest was identified using Spearman correlation.
